# Preventing plasmid multimer formation in commonly used synthetic biology plasmids

**DOI:** 10.1101/2024.07.23.604805

**Authors:** Elizabeth Vaisbourd, Anat Bren, Uri Alon, David S. Glass

**Affiliations:** Department of Molecular Cell Biology, Weizmann Institute of Science, Rehovot, Israel 76100

**Keywords:** plasmids, multimers, concatemers, recombination, nanopore sequencing, long-read sequencing

## Abstract

Plasmids are an essential tool for basic research and biotechnology applications. To optimize plasmid-based circuits, it is crucial to control plasmid integrity, including the formation of plasmid multimers. Multimers are tandem repeats of entire plasmids formed during replication by failed dimer resolution. Multimers can affect the behavior of synthetic circuits, especially ones that include DNA-editing enzymes. However, occurrence of multimers is not commonly assayed. Here we survey four commonly used plasmid backbones for occurrence of multimers in cloning (JM109) and wild-type (MG1655) strains. We find that multimers occur appreciably only in MG1655, with the fraction of plasmids existing as multimers increasing with both plasmid copy number and culture passaging. In contrast, introduction of multimers into JM109 can produce strains containing only multimers. We present an MG1655 *ΔrecA* single-locus knockout that avoids multimer production. These results can aid synthetic biologists in improving design and reliability of plasmid-based circuits.

## Introduction

Plasmids have been a crucial tool in engineering living systems for decades, in particular for engineering bacteria ^1–7^. Plasmids are small, circular DNA molecules that replicate independently of the bacterial chromosome, and are used for cloning, protein expression and genetic circuit assembly ^8,9^. The use of plasmids in synthetic biology allows for easy genetic manipulation ^10^, as compared to genetic engineering of chromosomes ^11^.

During plasmid replication, errant segregation may occur, leading to the formation of plasmid tandem repeats called multimers or concatemers ^12–15^. These multimers tend to accumulate over time and increase in number of repeats per molecule, a process known as the dimer catastrophe (Fig. 1A) ^16–18^. In bacteria, the presence of multimers with multiple origins of replication per molecule causes a decrease in the number of plasmid DNA molecules while maintaining the number of origins of replication ^16^. This affects the way in which plasmids are partitioned into daughter cells. For example, in randomly partitioned plasmids (e.g., with pColE1 origins), each daughter receives a random selection of molecules, not origins of replication ^16,19,20^. This random partition statistically increases the chance of daughter cells inheriting no plasmids at all ^19,20^. Additionally, the large DNA molecules increase the metabolic burden on the bacteria ^9^. Although molecular events leading to multimer formation and resolution have been studied ^12,15,21^, it is not clear to what extent multimers form during routine use in commonly used plasmids and laboratory strains.

**Figure 1.**
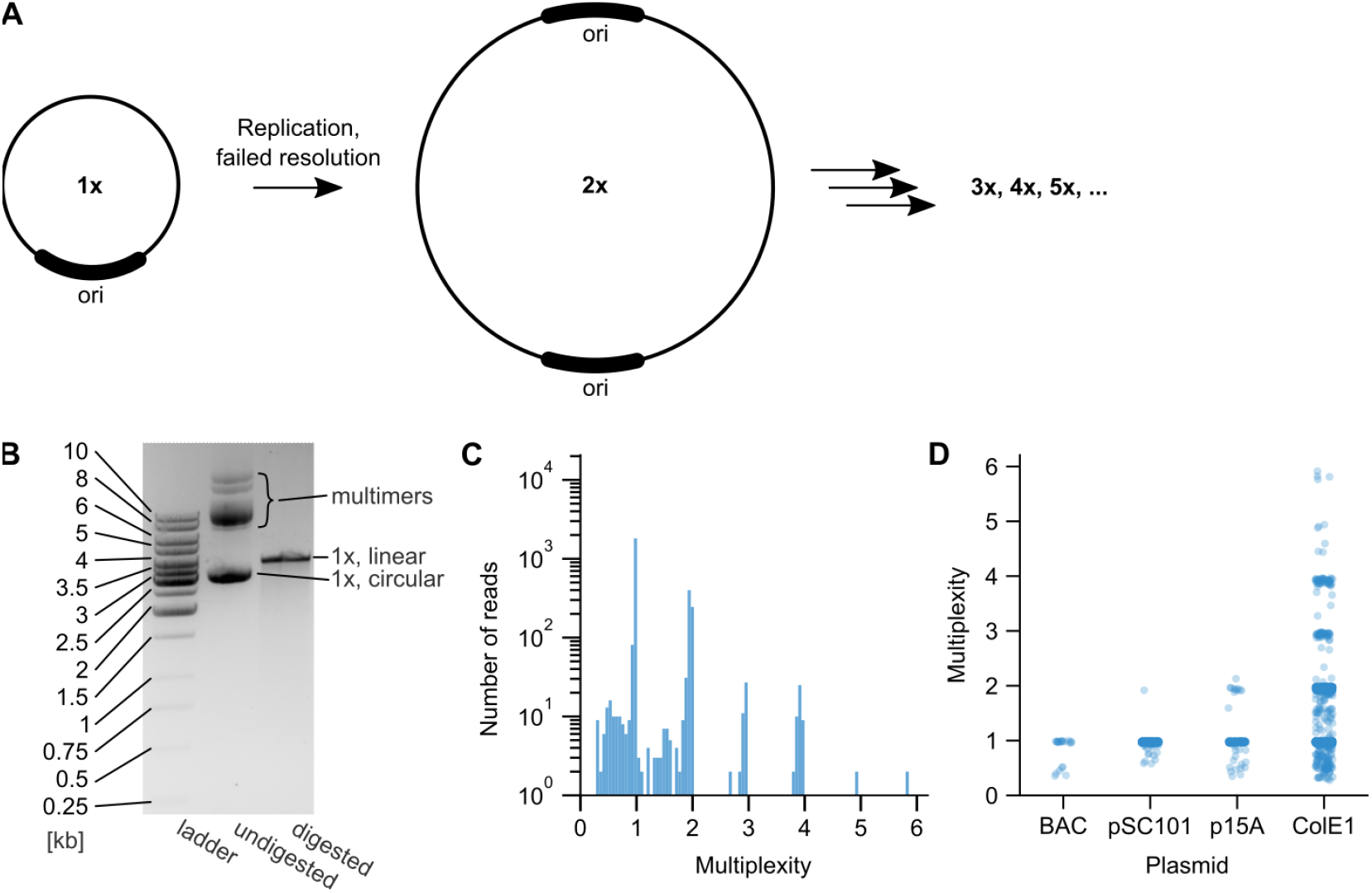
Long-read sequencing technology enables reliable detection of plasmid multimers. A. Schematic representation of multimer formation. During DNA duplication, failed resolution of plasmid dimers leads to persistence of multimers. Subsequent failures of resolution create trimers, tetramers, etc. B. Gel electrophoresis of the ColE1 plasmid isolated from MG1655 grown from a glycerol stock. Alongside a standard ladder (left lane), the plasmid sample was run as-is (center lane), and linearized by digestion (right lane) with SpeI, which cuts once in the singlet plasmid. Expected singlet (“1x”) length is 3660 bp. Note that the undigested plasmid may include forms such as supercoiled and nicked which can be hard to distinguish from multimers (see text). C. Histogram of mapped read lengths of the plasmid in (B) sequenced by nanopore sequencing. “Multiplexity” is the read length normalized by the plasmid’s nominal singlet (“1x”) length. Note that the longest read is a hexamer over 20kb. D. Distributions of mapped read lengths for all four plasmids in MG1655.

The presence of multimers can confound experiments. For example, mutations can occur in one of multiple repeats of a multimer plasmid but inherited with a wild type by physical linkage, thus changing plasmid behavior and selective pressure. Such unpredictability can be even greater for synthetic circuits utilizing DNA-editing enzymes such as recombinases ^8,22,23^ which could cut within or between multimers on the same DNA molecule. Plasmid integrity and stability are thus important to track both in synthetic circuit performance and when performing competition and evolution experiments ^8,24^.

### Factors affecting multimer formation

One aspect likely to influence the prevalence of multimers is the plasmid replicon ^12,14,25^, the genetic element that regulates the rate and timing of plasmid replication ^15,16,19^. Replicons control plasmid copy number in various ways, resulting in copy numbers that can range between 1 and several hundred ^26^. Copy numbers of plasmids have been quantified by qPCR ^20^ and microscopy ^19,27^. The origins and associated control elements of the replicon determine whether plasmids segregate into daughter cells randomly (e.g., with ColE1 origin) ^19^ or in a regulated fashion (e.g, F plasmids) ^25^. As part of the replication machinery, the origin and control elements also contain multimer resolution systems ^13^.

Likewise, the chromosomal background of a bacterial strain can affect multimer formation ^12^. Multimers are plasmids harboring two or more repeats of the same plasmid in tandem, that are caused by homologous recombination followed by a failed segregation site-specific recombination ^12,21^. The formation of multimers depends on chromosomal recombinases, namely recA, as well as recB, recC and recF ^12^.

### Detection of multimers

The prevalence of plasmid multimers has been traditionally assayed by gel electrophoresis ^28,29^. Gel electrophoresis, however, can be difficult to interpret. Even verified singlet plasmid samples can present on a gel as several bands, depending on the topology of the DNA ^28,29^. Linearized, circular, supercoiled, and nicked plasmids all run through agarose at slightly different speeds (Fig1B) ^14,30^. Owing to their larger sizes, multimers run slower than monomers in gel ^12^, but need to be distinguished from other plasmid forms in undigested plasmid isolates (Fig 1B) ^14^. Restriction digestion assays, frequently used to verify plasmid identity and sequence, do not faithfully reveal information about the plasmid length. Multimers, once digested, result in fragments of the same length as digested singlet plasmids (Fig 1B). Thus, barring structural variants created during multimerization, gel electrophoresis of digested plasmids will not indicate the presence of multimers.

It is also difficult to assay multimers using short-read sequencing. In theory, Sanger sequencing can detect structural variants resulting from faulty resolution of plasmid replication which may accompany some multimer formation. Because Sanger sequencing primarily detects consensus sequence, this requires that the structural variant dominates the sample. It also requires a primer that happens to be next to the variant breakpoint. High-throughput sequencing based on short reads could find such variants at sufficiently high read depths even if they do not dominate the sample, but they still do not reveal whole-plasmid tandem repeats without other structural variation.

The advent of long-read technologies, such as nanopore sequencing ^31,32^, provides a solution to the limitations of gels and short-read sequencing. By sequencing entire plasmids, multimers are directly apparent in histograms of read lengths (Fig. 1C-D).

### Multimer formation in common synthetic biology plasmids and strains

In this study, we examine plasmid multimerization dynamics in two common *Escherichia coli* strains: the cloning strain JM109 and the wild-type MG1655. We survey multimers in four common plasmid types based on BioBrick vectors, with pColE1 (BioBrick pSB1XX), p15A (BioBrick pSB3XX), pSC101 (BioBrick pSB4XX), and bacterial artificial chromosome (BAC, derived from the F plasmid) origins of replication ^26^. We measured multimers using nanopore sequencing and found that multimers increase with copy number in the wild-type strain MG1655, but not in the cloning strain JM109. The prevalence of these multimers in MG1655 increased with additional passaging of the strain. We also produced an MG1655 *ΔrecA* knock-out which shows little multimer formation, as in JM109. This information will aid in improved design, reliability, and interpretation of synthetic circuits.

## Results

In this study, we sought to quantify multimer dynamics in plasmids along a typical “life cycle” of usage (Fig. 2A). To this end, we assayed plasmids widely used in synthetic biology at three timepoints: before transformation, after transformation and growth from a colony, and after growth from a glycerol stock. This included amplification growth in a conventional cloning strain, JM109, and in the wild-type strain MG1655 ^8^. To account for biases that might stem from differences in origin of replication, we used plasmids spanning a range of copy numbers ^26^: bacterial artificial chromosome (BAC, ∼1 copy), pSC101 (“low-copy,” ∼5 copies, BioBrick backbone pSB4A3), p15A (“medium-copy,” ∼10-12 copies, BioBrick backbone pSB3A3), and ColE1 (“high-copy,” ∼500-700 copies, BioBrick backbone pSB1C3). We quantified multimer distributions using long-read nanopore sequencing of the BAC and plasmids.

**Fig. 2.**
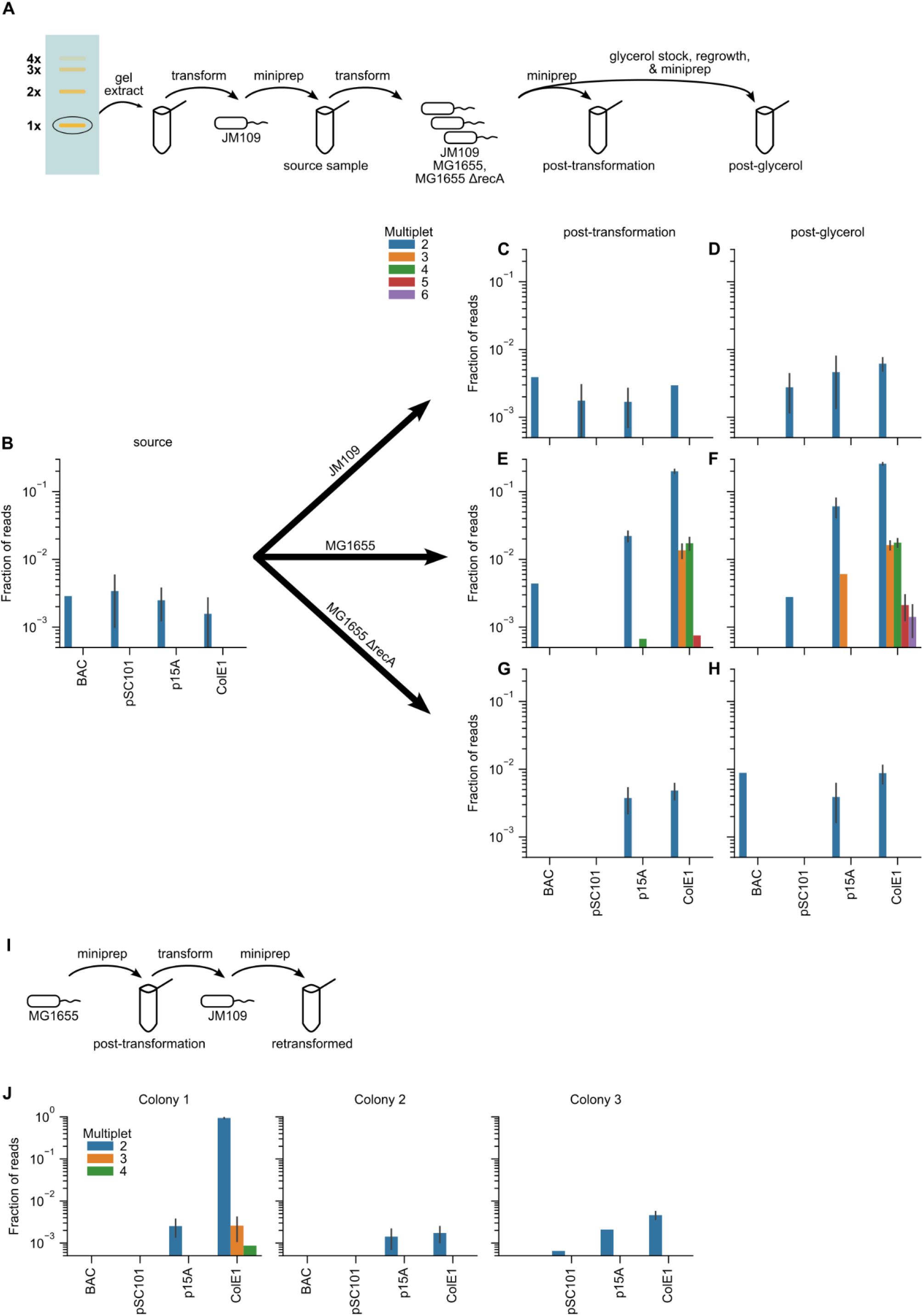
Fraction of plasmids existing as multimers increases with copy number, but are mostly eliminated by recA deletion, whereas transformation of multimers formed in a *recA*^*+*^ strain can lead to stable multimers in a ΔrecA strain. A. Schematic of plasmid isolation and transformations, yielding initial, nominally singlet “source” samples, followed by “post-transformation” and “post-glycerol” samples. B. Distributions of multimer states of plasmids in the source samples (nominally singlet from JM109). C-H. Distribution of multimer states of plasmids post-transformation (C,E,G) and post-glycerol (D,F,H) for JM109 (C,D), MG1655 (E,F), and MG1655 ΔrecA (G,H). I. Schematic of retransformation protocol of MG1655 “post-transformation” (A, E) samples back into JM109. Three colonies of plasmids were sequenced from the retransformation except for the BAC sample, for which the transformation yielded only 2 colonies. J. Distributions of multimer states of retransformed samples. Note that the ColE1 plasmid yielded nearly 100% dimer in 1 out of the 3 colonies. Error bars represent counting error based on the number of reads. Absence of error bars indicate that the measurement represents a single sequencing read.

We initiated the screen by isolating singlet plasmids. To do so, we size-separated plasmids via gel electrophoresis and extracted singlet-length bands for each plasmid. To obtain sufficient material for experimentation, we transformed these gel extracts into JM109 and isolated plasmid DNA from the transformed strains. We refer to these samples as “source” samples, because they simulate a plasmid sample one might expect to receive from an outside lab. To verify the singlet status of the presumed singlet plasmids, we sequenced these plasmids via nanopore sequencing. We found that these contained up to ∼0.3% dimers (Fig. 2B).

To test the stability of the singlets in JM109, we transformed the isolated singlet plasmids again into JM109. For each plasmid, we picked a single colony, which we then grew in overnight culture. We isolated plasmid DNA directly from this transformed culture. We refer to these samples as “post-transformation.” We also saved the culture as a glycerol stock, which we then regrew in a new overnight culture. We isolated plasmid DNA from this second culture, which we referred to as “post-glycerol.” Such a culture represents what would normally be used for experiments on the strain containing the plasmid. In both the post-transformation and post-glycerol samples, we found that less than ∼0.6% of total plasmid content appeared as dimers, and we observed no multimers with more than two tandem repeats (Fig. 2C-D).

To examine the dynamics of plasmid multimers in the widely used wild type MG1655 strain, we transformed the source samples (Fig. 2B) into MG1655. As with JM109, we isolated plasmid DNA from both an initial post-transformation overnight and from culture grown from glycerol. In the post-transformation sample (Fig. 2E), less than ∼0.4% of the BAC and low-copy plasmids were dimers. The medium-copy plasmid had ∼2% dimers and <∼0.06% higher-order multimers. Of the high-copy plasmid ColE1, 20% were dimers, 1.3% trimers, 1.7% tetramers, and <∼0.075% higher-order. In the post-glycerol samples (Fig. 2F), the fraction of plasmids existing as multimers increased overall, reaching 6% dimers in medium copy p15A and 25% dimers in high-copy ColE1. Hexamers were observed in the high-copy plasmid at ∼0.15%.

We next sought to test whether multimer formation and maintenance can be abolished in MG1655. To this end, we knocked out *recA* from MG1655 (Methods), as *recA* is knocked out in JM109 and is known to be necessary for multimer formation ^12^. In this strain (Fig. 2G-H), dimers were present in percentages less than ∼0.4% - 0.8% across all conditions, with no higher order multimers, similar to the results JM109.

Finally, we tested to see whether retransformation of multimeric plasmids into JM109 can increase the fraction of multimer plasmids even in the cloning strain (Fig. 2I-J). We transformed the MG1655 “post-transformation” samples into JM109 and isolated plasmid DNA from 2-3 colonies for each plasmid type (Fig. 2I). We found that for the high-copy plasmid, 1 of 3 colonies yielded nearly 100% dimers, indicating that the dimer form, if transformed into JM109, is stable and does not get resolved into singlet plasmids (Fig. 2J).

## Discussion

In this study, we used nanopore sequencing to quantify multimer formation in common synthetic biology plasmids and strains following a typical “plasmid life cycle” through transformation, culture growth, and regrowth. We found that in the cloning strain JM109, the only multimeter formed at detectable levels are dimers, and these are maintained below 1% of the plasmid sample. In the wild type MG1655, multimers increased with copy number and passaging of the culture through an additional overnight culture. We found up to six tandem repeats of the singlet sequence, with up to ∼30% multimers of all degrees. Upon knockout of *recA* in MG1655, multimer formation returned to basal JM109 levels. We found that transformation of multimeric plasmids into JM109 can lead to cultures with no singlet molecules (100% dimers and higher order multimers).

These results l are intended as a brief survey, and have a number of limitations. First, the “source” plasmid samples contained ∼0.3% dimers. It is possible that these influence the formation of multimers in the ensuing experiments. However, given the relatively constant percentage in JM109 and MG1655 *ΔrecA* strains, these may represent a natural fraction of plasmids undergoing replication at the moment of isolation. Second, the lengths, antibiotic resistance markers, and insert sequences varied across the plasmids tested (Methods). The metabolic burden imposed by these plasmids may thus differ, possibly affecting the exact multimer fractions.

Controlling plasmid sequences is crucial for proper understanding of plasmid function. We hope that the quantification of plasmid multimers in this work will be helpful for future experiments. Likewise, we hope that the *recA* knockout strain will provide a stable wild type-like chassis for engineering plasmid-based strains without multimers.

## Methods

### Strains and plasmid construction

JM109 was sourced from RBC (Cat #RH718), MG1655 from the Coli Genetic Stock Center (CGSC #6300). To prepare the *ΔrecA::KanR* strain, the knockout region was transferred to MG1655 (CGSC #6300) from the Keio collection strain JW2669 (BW25113 *ΔrecA::KanR*) using P1 phage transduction ^33^. The high-copy plasmid was constructed via Gibson assembly by moving the insert of pNR230 ^34^ to a pSB1C3 backbone. Other plasmids were previously published (see Table 1).

**Table 1:** details on the plasmids used in this study.

| Plasmid | Origin | BioBrick backbone | Copy number | Resistance marker | Length (bp) | Source |
| --- | --- | --- | --- | --- | --- | --- |
| pDSG602 | BAC |  | ~1 | Chloramphenicol | 10323 | <sup>8</sup> |
| pDSG467 | pSC101 | pSB4A3 | ~5 (“low”) | Ampicillin | 5958 | <sup>8</sup> |
| pDSG588 | p15A | pSB3A3 | ~10-12 (“medium”) | Ampicillin | 4740 | <sup>8</sup> |
| pDSG444 | ColE1 | pSB1C3 | ~500-700 (“high”) | Chloramphenicol | 3660 | This work |

### Growth conditions

Cultures were all grown overnight in 3 mL LB, shaken at 300 rpm at 37C.

### Gel electrophoresis

Samples were run on 1% agarose gels at 100V for ∼2 h.

### Plasmid isolation

Plasmid isolations were all performed using Qiagen Qiaprep spin miniprep kit (Cat# 27104), including the PB step.

### Plasmid digestion

Plasmid digestion for Fig. 1B was performed using NEB SpeI (Cat #R1033) following manufacturer’s recommended protocol. Digestion was run for 2 h.

### Nanopore sequencing and analysis

Nanopore sequencing was all performed by Plasmidsaurus, following their standard plasmid sequencing service. Sequences were aligned to known plasmid references using minimap2 2.21-r1071 and processed using samtools 1.12 and seqkit 2.2.0. Subsequent analysis was performed in python 3.9.7 using Bipython 1.79 and pysam 0.16.0.1. Total aligned overlap with reference was collected using pysam. To obtain “multiplexity” values, overlaps were divided by the singlet plasmid length (see Fig. 1D). For Figs. 2-3, these multiplexity values were rounded up to the nearest integer, which assumes that fragments that cover, for example, 1.5 singlet lengths come from the shortest possible multimer.

## Acknowledgements

The authors would like to thank the Alon lab for helpful feedback and comments on the manuscript. Funding was provided by the European Research Council (ERC) under the European Union’s Horizon 2020 research and innovation program (grant agreement no. 856587) and by the Israel Science Foundation (grant agreement no. 1966/22). D.S.G. was funded as a member of the Zuckerman Postdoctoral Scholars Program. U.A. is the incumbent of the Abisch-Frenkel Professional Chair.

